# Protective effects of nicotinamide in a mouse model of glaucoma DBA/2 studied by second-harmonic generation microscopy

**DOI:** 10.1101/2024.03.07.583928

**Authors:** Vinessia Boodram, Hyungsik Lim

**Affiliations:** Department of Physics and Astronomy, Hunter College of the City University of New York, New York, NY 10065; School of Optometry, Indiana University, Bloomington, IN 47405

## Abstract

Glaucoma is a blinding disease where the retinal ganglion cells and their axons degenerate. Degradation of axonal microtubules is thought to play a critical role in the pathogenesis, but the mechanism is unknown. Here we investigate whether microtubule disruption in glaucoma can be alleviated by metabolic rescue. The morphology and integrity of microtubules of the retinal nerve fibers were evaluated by second-harmonic generation microscopy in a mouse model of glaucoma, DBA/2, which received a dietary supplement of nicotinamide to reduce metabolic stress. It was compared with control DBA/2, which did not receive nicotinamide, and non-glaucomatous DBA/2-*Gpnmb+*. We found that morphology but not microtubules are significantly protected by nicotinamide. Furthermore, from co-registered images of second-harmonic generation and immunofluorescence, it was determined that microtubule deficit was not due to a shortage of tubulins. Microtubule deficit colocalized with the sectors in which the retinal ganglion cells were disconnected from the brain, indicating that microtubule disruption is associated with axonal transport deficit in glaucoma. Together, our data suggests significant role axonal microtubules play in glaucomatous degeneration, offering a new opportunity for neuroprotection.

## 1. Introduction

Glaucoma is a leading cause of blindness worldwide (1, 2), where nearly half of the retinal ganglion cells (RGCs) and their axons are irreversibly lost by the time of diagnosis. The risk factors include aging and high intraocular pressure (IOP), but the pathogenic mechanism is poorly understood hampering the prevention of vision loss. Axonal microtubules have been postulated to play a crucial role in the disease (3-5). It has been demonstrated that in DBA/2 mice (DBA), a well-characterized model of inherited glaucoma (6-9), axonal microtubules decay with age more rapidly than the RGC axons (i.e., the retinal nerve fibers) (10). The degradation of microtubules would impair axonal transport depriving the RGCs of essential tropic and metabolic supports. One of the possible drivers of microtubule loss is dysfunctional cellular metabolism, which could compromise active regulations stabilizing the microtubule assembly. Microtubule stability and metabolic equilibrium are co-dependent: Conversely, the disturbance of microtubules, which enable intracellular transport of mitochondria, could induce local energetic shortfall in the distal compartments. Lowering metabolic stress with metabolites, e.g., nicotinamide adenine dinucleotide (NAD) or its precursor nicotinamide (NAM), has been shown to delay or halt axonal degeneration (11) and protect RGCs against glaucoma (12, 13). It is plausible that the mechanism of neuroprotection by NAD/NAM involves axonal microtubules.

Here, using NAM as a paradigm, we investigate how axonal microtubules of RGCs respond to metabolic rescue. Second-harmonic generation (SHG) microscopy was employed for measuring microtubules in the retina nerve fibers, which, by virtue of the sensitivity to uniformly polarized microtubules (14-16), provides distinctive information regarding the cytoskeleton’s integrity.

## 2. Results

An overview of the experiment is shown in Fig 1(a). A group of DBA mice (N=41, 11 from 8 males and 30 from 16 females) received a dietary supplement of NAM prophylactically from 6 months of age at a low dose of 550 mg of body weight per day, as previously demonstrated (12, 13). As a control, another group of DBA mice received a normal diet without NAM at the same ages (N=33, 13 from 8 males and 20 from 12 females). The strain-matched, homozygous *Gpnmb+* mice (DBA*-Gpnmb+*) were used as non-glaucomatous control (17) (N=19, 13 from 8 males and 6 from 4 females). At a desired age, the IOP was measured using a TonoLab tonometer (18) on three different dates prior to imaging, and the average was taken. Mosaics were acquired over a region around the optic nerve head. Image processing was carried out to normalize the SHG intensity and evaluate the thickness of the retinal nerve fibers (10). The normalized SHG intensity divided by thickness, namely SHG density, provides a measure of the density of axonal microtubules at a microscopic resolution. The mean SHG density was evaluated over the retinal nerve fibers in the mosaic. Also, the thickness was integrated over the region to yield the volume of the retinal nerve fibers.

**Fig 1.**
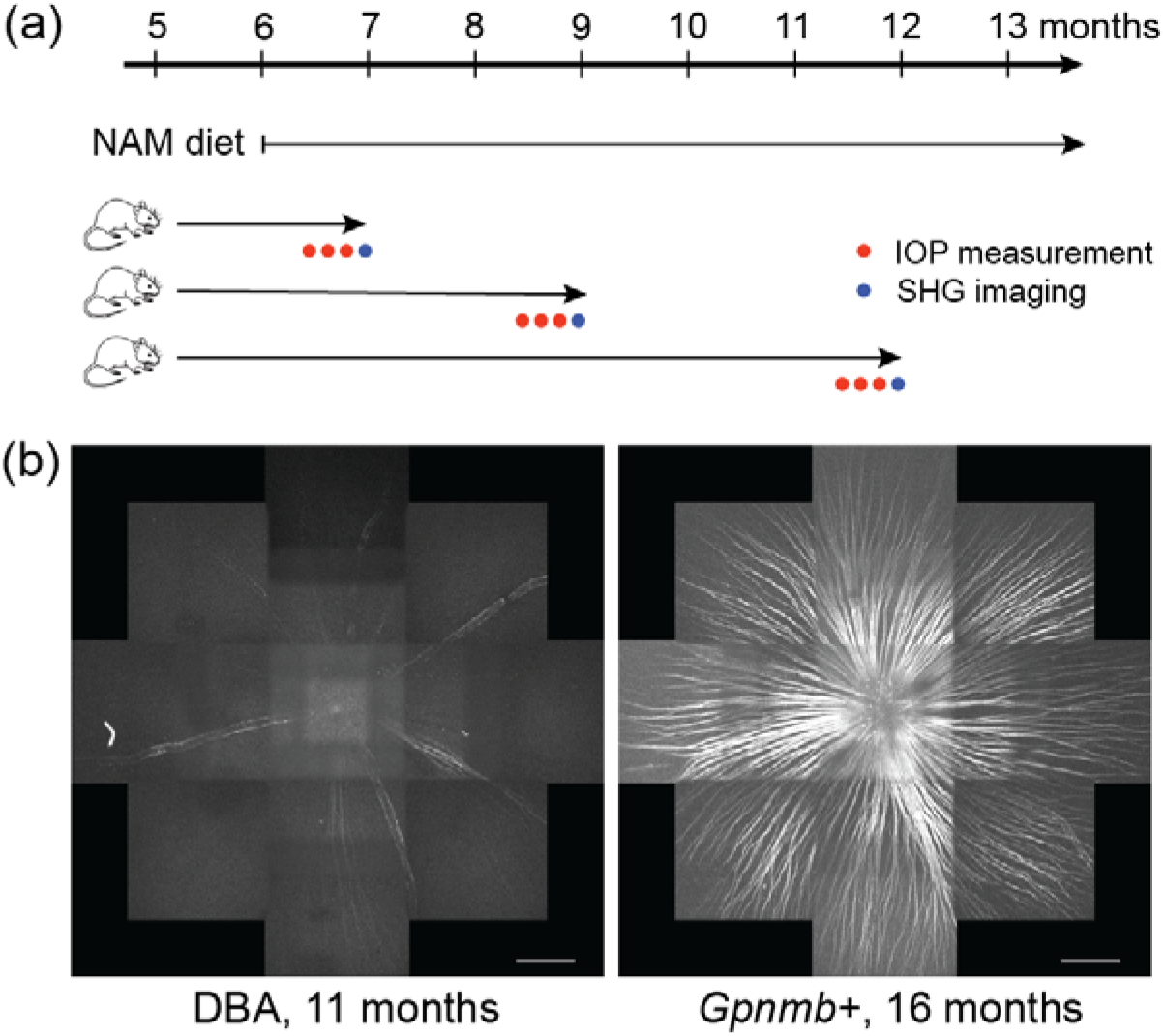
Overview of experiment. (a) Timeline of measurements. (b) Degenerated DBA and healthy *Gpnmb+* retinas visualized by SHG. Scale bars, 300 μm.

### NAM improves morphology more effectively than microtubules of the retinal nerve fibers

The effect of NAM diet was investigated on three properties of the eye (i.e., IOP, volume, and mean SHG density) (Fig 2). Overall, the variations of DBA data were greater than those of *Gpnmb+* representing primarily the experimental errors. The age-dependent changes of the DBA eyes were analyzed using a multiple linear regression model containing three main effects that are known to influence the pathology (i.e., age, sex, and diet) as well as two interactions (summarized in Table 1). While the IOP of *Gpnmb+* mice remained stable, that of DBAs without NAM diet increased with age, as expected. In DBA mice, the rate of IOP elevation with age varied depending on diet, which was significantly lower with NAM than without NAM (*p*=.046). Diminished IOP elevation has also been observed in the prior study but at a higher dose of NAM (12, 13). Interestingly, we found that the benefit of NAM supplement was significant for protecting the morphology, but not microtubules, of the retinal nerve fibers. The volume of the retinal nerve fibers in DBA was protected against age-dependent loss with NAM diet (*p*=.049). While the mean SHG density also decayed more slowly with NAM diet, it was not statistically significant (*p*=.43).

**Table 1.** Summary of multiple linear regression model, *y*= *α* + *β*_*age*_ · *Age* + *β*_*sex*_ · *Sex* + *β*_*diet*_ · *Diet* + *β*_*intl*_ · *Age* · *Sex* + *β*_*int2*_ · *Age* · *Diet* + *ε*.

|  | Dependent variables |  |  |
| --- | --- | --- | --- |
|  | IOP (mmHg) | Volume (x10<sup>6</sup> μm<sup>3</sup>) | Mean SHG density (a. u.) |
| Age | 0.60***<br>(0.12) | -1.50***<br>(0.39) | -25.04***<br>(8.26) |
| Sex | -1.98<br>(1.56) | -4.09<br>(5.08) | -87.27<br>(106.52) |
| Diet | 2.14<br>(1.48) | -7.80<br>(4.83) | -66.32<br>(101.30) |
| Age·Sex | 0.17<br>(0.15) | 0.58<br>(0.49) | 9.04<br>(10.33) |
| Age·Diet | -0.29**<br>(0.14) | 0.94**<br>(0.47) | 7.86<br>(9.84) |
| Constant | 5.24***<br>(1.22) | 20.76***<br>(3.98) | 373.52***<br>(83.49) |
| Observations | 74 | 74 | 74 |
| R<sup>2</sup> | 0.47 | 0.25 | 0.18 |
| Adjusted R<sup>2</sup> | 0.43 | 0.20 | 0.12 |
| F Statistic<br>(df = 5; 68) | 12.03*** | 4.61*** | 2.93** |

Note: \* $p < .1$ ; \*\* $p < .05$ ; \*\*\* $p < .01$

**Fig 2.**
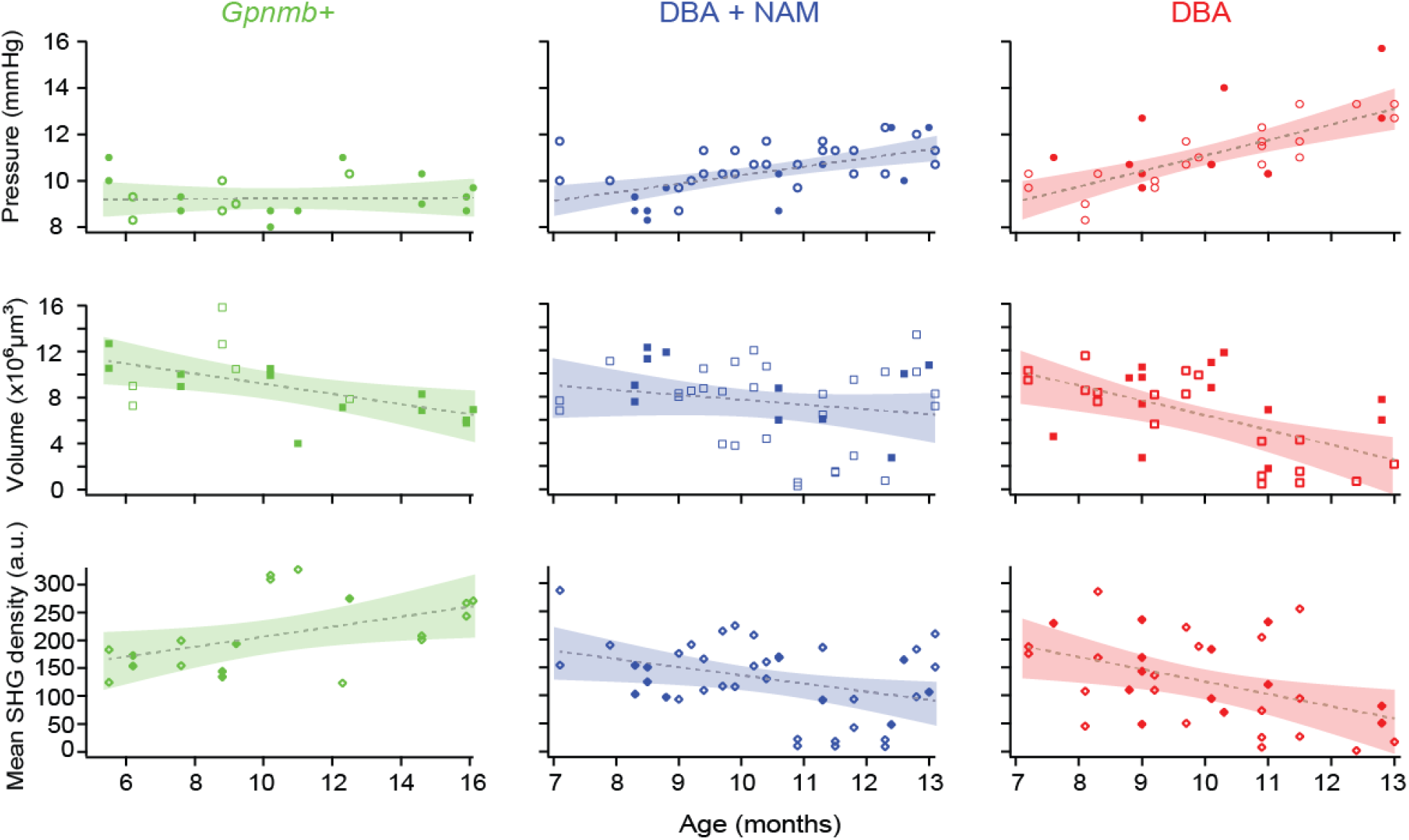
Age-dependent effects of NAM dietary supplement on the DBA retinas. The IOP, the volume and mean SHG density of the retinal nerve fibers are shown for three groups, i.e., *Gpnmb+*, DBA with and without NAM diet. Solid and open markers are male and female, respectively. Dashed lines are simple linear regressions with age and shaded bands are 95% confidence bounds.

### The mean SHG density and volume are negatively correlated with IOP

In addition to age-dependent changes, we investigated how the SHG-derived parameters are related to the IOP, another major risk factor of glaucoma (Fig 3). In DBA mice without NAM diet, we found modest negative correlations at the level of significance between the IOP and the mean SHG density (Pearson correlation coefficient r=-0.42, *p*=.015) and the volume (r=-0.37, *p*=.033), implying that both axonal microtubules and RGC axons were lost possibly as downstream effects of the IOP elevation. Then we examined how the dependence on IOP was affected by the NAM diet. Specifically, we asked whether the protective effect of NAM was, instead of being a direct consequence of bolstered metabolism, mediated entirely by the IOP reduction, which would preserve the correlation coefficient. On the other hand, if NAM had an additional mode of action to prevent the loss of RGC axons, the correlation was expected to decrease as the IOP explains less of the variation. Unfortunately, the correlation coefficient between the volume and the IOP could not be determined with confidence for the group of DBA mice with NAM diet (r=-0.19, p=.24), leaving the inquiry unresolved.

**Fig 3.**
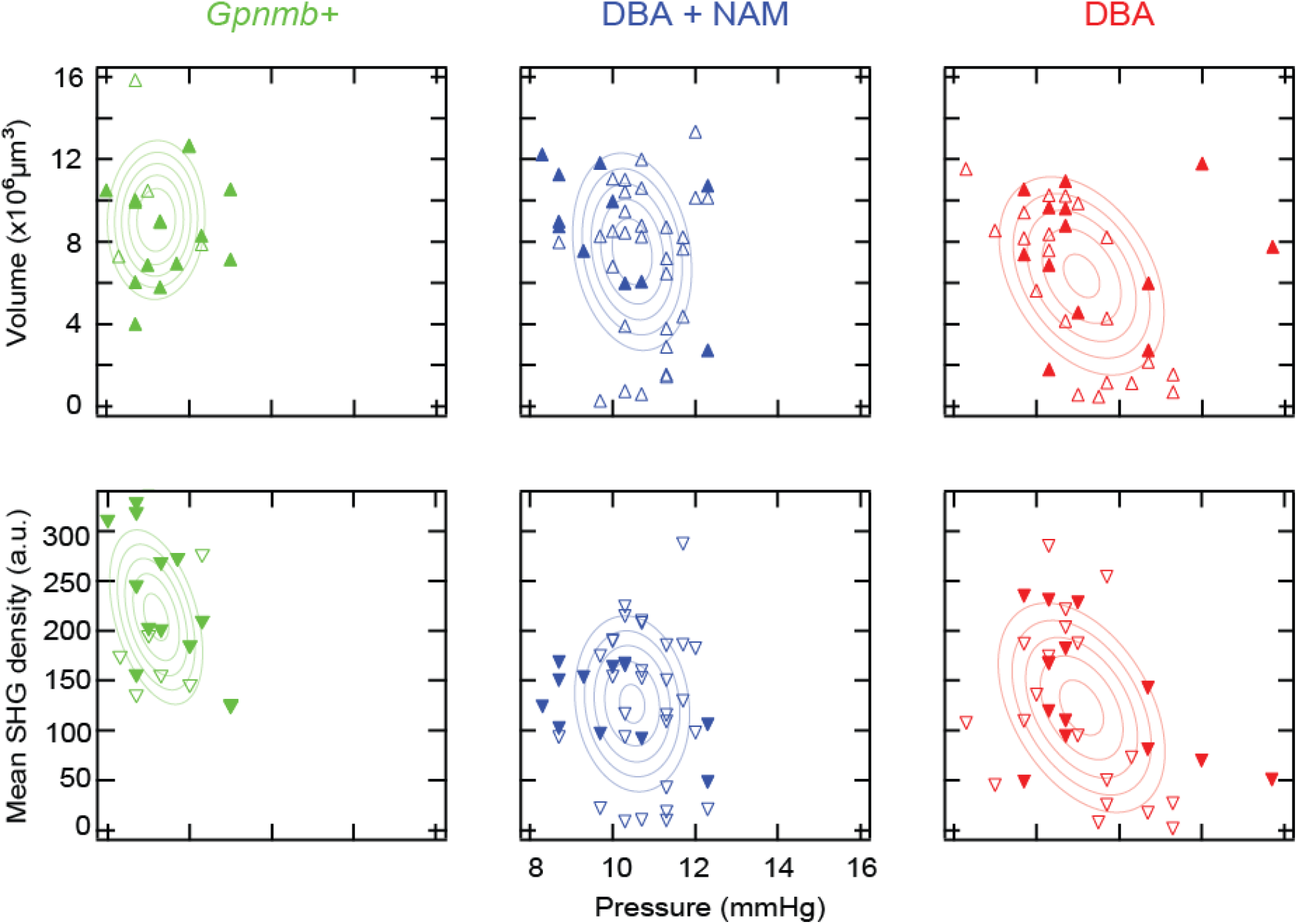
Correlation between morphology and microtubules versus IOP for three groups. Solid and open markers are male and female, respectively, overlaid with the best fits to the bivariate normal distribution.

### Microtubule deficit evolves in a NAM-dependent manner during glaucoma progression

Axonal microtubules are disrupted before the loss of RGC axons during glaucomatous degeneration leading to a pathological state called *microtubule deficit* (10), where axonal microtubules exist in lower quantities than normal for the caliber. Due to the differential decay rates of axonal microtubules and morphology in response to NAM diet, it was anticipated that microtubule deficit would follow different trajectories during glaucoma progression. To illustrate this, the mean SHG density and volume were analyzed jointly in the same retinas. Provided that the rate coefficients of the mean SHG density ρ and volume V are given by α and β, respectively,

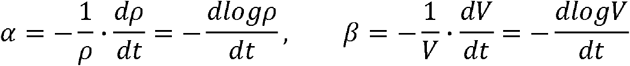

then the ratio of coefficients is obtained from the slope of a log-log plot.

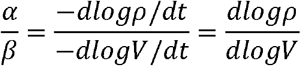

Microtubule deficit results for a ratio greater than unity *(α⁄ β* > 1), as in the case that the volume is relatively protected, i.e., the loss of volume is slower than microtubule disruption. The relative decays were compared for two populations of DBA retinas with and without NAM diet (Fig 4). An evident pattern was exhibited where samples of the group with NAM diet were shifted toward microtubule deficit compared to the group without NAM diet, confirming the decay of volume was dampened relative to that of axonal microtubules in those samples. Furthermore, the DBA retinas had higher probabilities of microtubule deficit. The result reproduces the previous findings (10) with different experimental protocols and apparatus, confirming microtubule deficit as a molecular marker of glaucoma.

**Fig 4.**
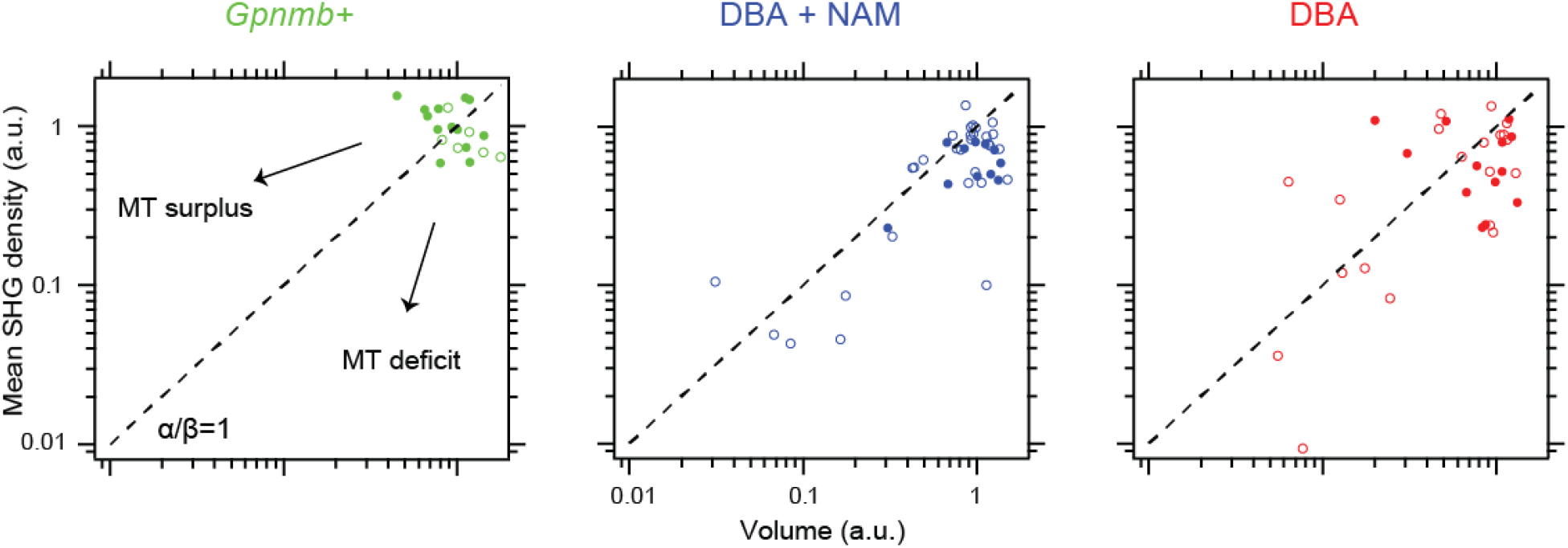
The relative course of degeneration of morphology and microtubules MT; microtubule. The parameters are normalized to the average values of *Gpnmb+* retinas. Solid and open markers are male and female, respectively. Dashed line, the unity slope.

### Microtubule deficit is not due to paucity of tubulins

Microtubule deficit could be either due to a shortage of tubulin monomers, e.g., from inadequate expression or transport to the RGC axons, or arising from a failure to stabilize the polymer, e.g., from dysfunctional regulation. To resolve these possibilities, we performed co-registration of immunofluorescence and SHG images. The DBA retinas were fixed immediately after SHG imaging and double immunostained against two types of cytoskeletal protein, i.e., class III beta-tubulin (βIII) and phosphorylated neurofilament (pNF). βIII is a neuron-specific isoform of tubulin, and pNF is normally enriched in axons but accumulates in the soma of degenerating RGC’s. To examine the microtubule integrity and the abundance of tubulins in the same regions, βIII immunofluorescence was co-registered with SHG images. A total of 12 DBA retinas were evaluated, primarily focusing on the age range between 10 and 12 months. In the retinas exhibiting pronounced sectorial degeneration (N=4), which is a characteristic in glaucoma pathology, the sectors of microtubule deficit always contained morphologically intact retinal nerve fibers (Fig 5(a), between dashed lines). Also, it also showed high signals of βIII tubulins inside the sector, i.e., no substantial impairment in the expression or transport. It can be therefore concluded that microtubule deficit is likely due to the instability of microtubules rather than a low level of tubulins.

**Fig 5.**
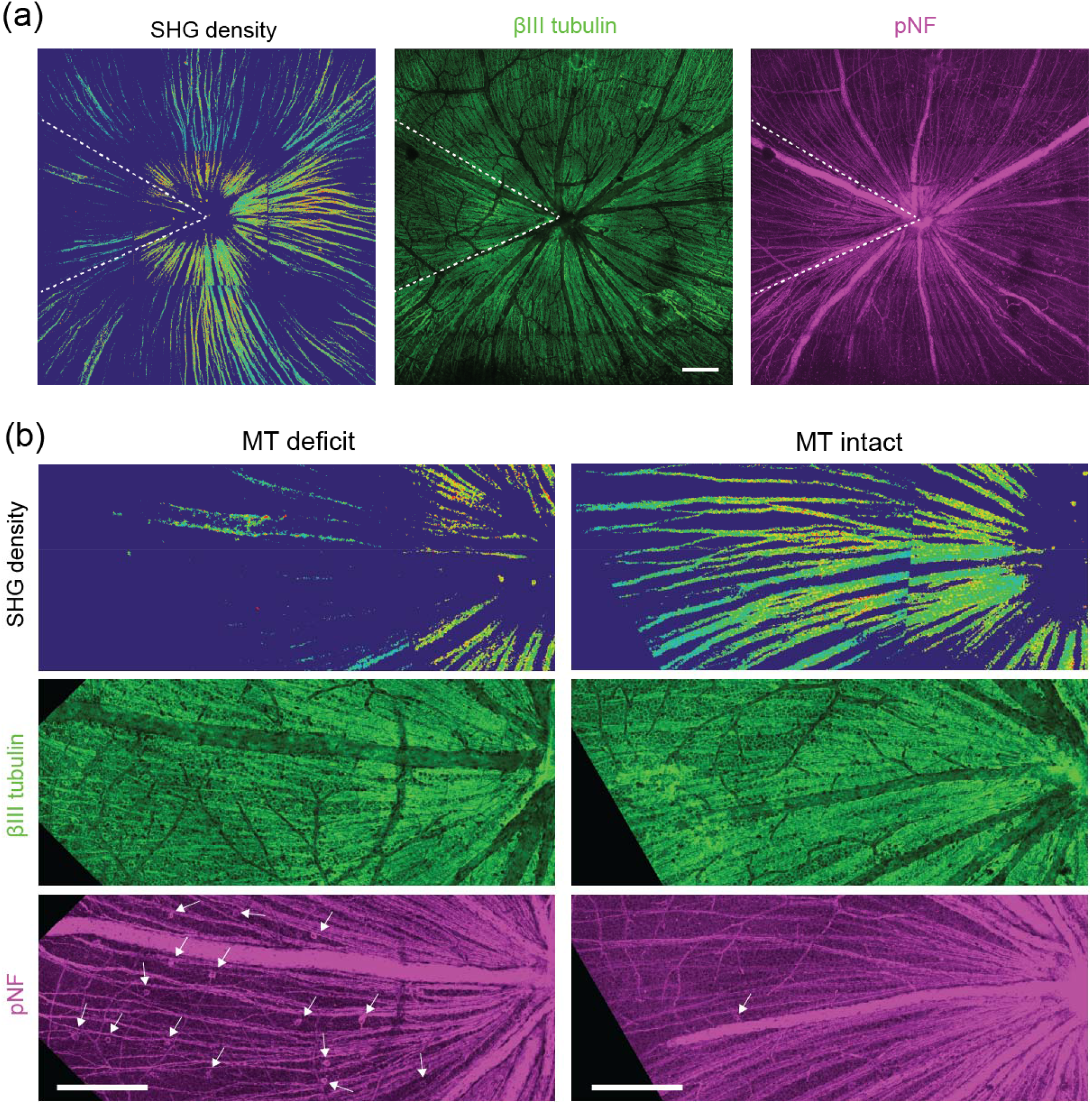
Microtubule deficit versus the distribution of cytoskeletal proteins, βIII tubulin and pNF. (a) Mosaics of a DBA retina (22 months age, female) by SHG and immunofluorescence against the cytoskeletal elements. (b) Comparing two sectors with deficient and intact microtubules. Arrows, pNF+ RGC somas. Scale bars, 200 μm.

### Loss of axonal microtubules colocalizes with the RGC’s disconnect from the brain

The somatic accumulation of pNF, which can be detected with monoclonal antibody lacking nonspecific staining of soma (2F11), is interpreted to suggest that the RGC is disconnected from the brain, as has been validated by retrograde tracing (19, 20). To investigate the relationship between microtubule and transport deficits, we compared SHG density with the distribution of pNF. We found that pNF+ somas represented a small subset of the RGC population (in the order of dozens in an area of approximately 1.8 mm diameter around the optic nerve head) (Fig 5(b)) and were absent in highly degenerated areas (Fig S1). Presumably, not all RGCs undergo this pathological fate during glaucomatous degeneration. The redistribution of pNF could be short-lived with pNF+ RGC somas disappearing altogether upon the cells’ death. Interestingly, pNF+ RGC somas were localized in sectors (Fig 5 and Fig S1) and routinely coincided with that of low SHG density (Fig 5(b)). The result, together with intact βIII signals, suggests that the RGC’s connection to the brain is likely to be severed at the time of microtubule disruption while the RGC axons are intact, i.e., at the stage of microtubule deficit.

## 3. Discussion

Taken together, our data supports a model in which NAM/NAD has several independent modes of action targeting distinct aspects across the pathways of glaucoma pathogenesis (Fig 6). The IOP elevation has a component that responds to NAM. The reduced ocular hypertension is partially responsible for protecting the retinal nerve fibers, but there also seems to be an IOP-independent pathway (‘?’ in Fig 6): NAM promotes the morphology of the retinal nerve fibers more significantly than microtubules. The differential response can be explained by theorizing that the loss of RGC axons is a compound effect of microtubule disruption and another necessary condition that is downregulated under metabolic stress and IOP-independent. Such an IOP-independent pathway is also consistent with the previous finding that NAM protects RGCs without altering the IOP (12, 13). Conceivably, even more facets of the RGC degeneration are modulated by NAM/NAD, as metabolism controls diverse cellular processes in most cell types. One of the challenges to unraveling the mechanism was unexplained variability of the DBA model, which should be addressable in future studies by employing IOP-induced rodent models (21). Detailed knowledge of the distinct actions of NAM can improve the precision of NAM-based glaucoma therapy.

**Fig 6.**
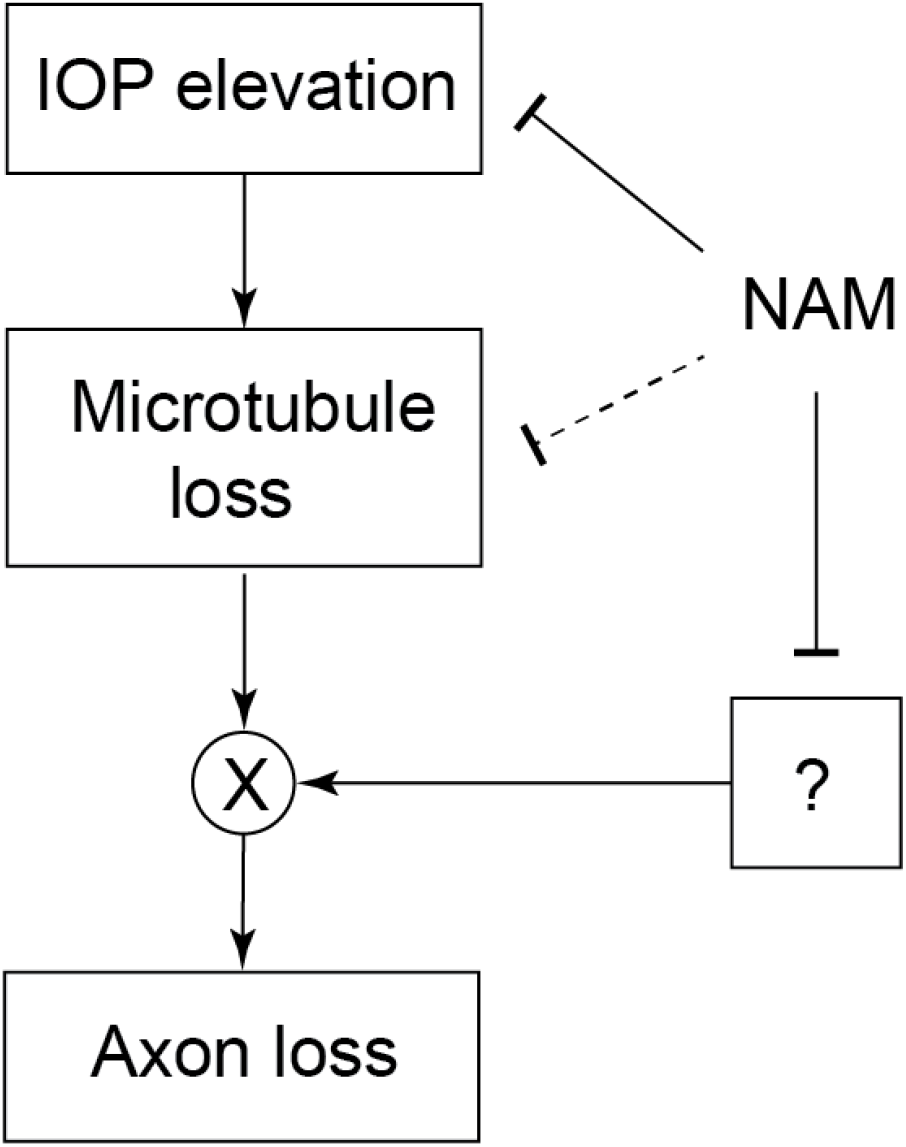
Model of the glaucomatous degeneration of RGC axons.

Axonal microtubules are involved in the glaucomatous RGC degeneration. We hypothesized that metabolic stress, such as mitochondrial abnormalities or depletion of metabolites, might destabilize axonal microtubules to induce microtubule deficit. The hypothesis predicts that microtubule deficit will be mitigated by NAM supplement. It is found that, despite having multiple mechanisms to protect RGC axons, NAM/NAD provides only limited protection of axonal microtubules. Microtubule disruption in glaucoma does not appear to be strongly related to metabolic decline. Instead, the primary pathogenic source could be the elevated IOP, which exhibits a negative correlation with axonal microtubules. It raises an important question regarding glaucoma therapy, i.e., whether the efficacy will be eventually hampered without reversing the degradation of axonal microtubules.

Critical insights have been gained into microtubule deficit. It has been shown that microtubule deficit is not a consequence of a shortage of tubulins in the RGC axon. Regarding whether deteriorating axonal microtubules play a causal role in the eventual loss of RGCs, our multimodal data reveals spatially overlapping, sectorial patterns of microtubule and axonal transport deficits. Correlated sectorial degenerations have allowed researchers to associate the loss of RGC axons with pathogenic insults at the optic nerve head (7, 22-24). Similarly, the alignment of axonal transport and microtubule deficits suggest that the cytoskeletal breakdown might be an intermediate stage in the progressive RGC death. The proposed role of axonal microtubules offers a new opportunity for neuroprotection against glaucomatous degeneration.

We have demonstrated that SHG imaging is well-suited for interrogating microtubule deficit. The information in the SHG signal is much distinguished from immunofluorescence against tubulin monomers, which is necessary but not sufficient for the integrity of microtubule assembly. Characterizing microtubule deficit across a large area of the retina is nontrivial with electron microscopy. By contrast, it can be surveyed readily across the whole retina by SHG imaging. Being sensitive to the functional form of axonal microtubules, SHG provides an ideal readout for dissecting the mechanistic role in glaucoma pathogenesis.

## Materials and Methods

### Animals

All procedures were approved by the Hunter College Institutional Animal Care and Use Committee (IACUC). DBA (DBA/2J, # 000671) and *Gpnmb+* (DBA/2J-*Gpnmb+*/SjJ, # 007048) mice were obtained from The Jackson Laboratory and housed in the animal facility at Hunter College.

### Pharmacology

Nicotinamide was given at a low dose of 550 mg of body weight per day(12, 13) by adding to standard pelleted chow (2750 mg/kg) (Bio-Serv).

### IOP measurement

IOP was measured using a TonoLab tonometer (Icare) while mice were awake (i.e., unanesthetized) and restricted. Three measurements were performed on consecutive dates prior to SHG imaging. The time of measurement was standardized at a fixed hour during daytime to avoid the intraday fluctuations (25).

### Tissue preparation

The animal was deeply anesthetized with isoflurane and the first eye was enucleated. After the enucleation of the second eye, the animal was euthanized by CO_2_ inhalation. The retinal flatmounts were prepared as previously described (10). Briefly, an incision was made along the corneal limbus, the lens and sclera were removed, and radial cuts were made to relieve the curvature. The flat-mounted retina was transferred to a glass bottom dish (MatTek Corp.) and incubated at room temperature in the Ames’ medium (A1420, Sigma-Aldrich) oxygenated with 95%O_2_/5%CO_2_.

### SHG microscopy

An experimental setup for SHG microscopy was similar to the previous study (10, 15). Briefly, 100-fs pulses at an 80-MHz repetition rate from a Ti:Sapphire laser (Chameleon Ultra, Coherent, Inc.) were used for the excitation. The output wavelength was 900 nm. The polarization state of excitation beam was controlled with half-and quarter-waveplates. A water-dipping microscope objective lens (HC FLUOTAR L 25x 0.95NA, Leica) was used to focus the excitation beam onto the sample. The average power was approximately 20 mW at the sample. The forward-propagating SHG from the sample was collected with an UV-transparent high-NA objective lens (UApo340 40× 1.35NA, Olympus), passed through a narrow-bandpass filter (<20-nm bandwidth) at a half of the excitation wavelength (400 nm), and then detected with a photomultiplier tube (PMT; H7422-40, Hamamatsu, Inc.). Images with 512×512 pixels were acquired, and the pixel dwell time was ∼ 3 μs. A region was imaged twice for orthogonal linear polarizations, which then were summed into a composite image. Z-stacks were acquired in a step of 2 μm. For creating mosaics, a total of 9 regions (742×742 μm^2^ each) were imaged on and around the optic nerve head at 1-mm radius.

### Immunohistochemistry and confocal microscopy

Immunohistochemistry was performed similar to the prior studies (20, 24, 26). After SHG imaging, the retinal flatmount was fixed with 4% paraformaldehyde for 20 minutes at room temperature. The sample was dehydrated sequentially in 25%, 50%, 75%, and 100% cold methanol for 15 min each and then permeabilized with dichloromethane for 2 hours. Then the retina was rehydrated in 75%, 50%, 25%, and 0% cold methanol for 15 min each. The sample was blocked in a buffer containing 5% normal serum and 0.3% Triton™ X-100 (Thermo Fisher Scientific, Inc.) for 3 hours. The sample was incubated in the primary antibody buffer at 4°C for 3 days. Rabbit and mouse monoclonal antibodies against βIII tubulin (EP1569Y, Abcam, Inc.) and pNF (2F11, EMD MilliporeSigma), respectively, were used at 1:300 dilution. The sample was incubated in the secondary antibody buffer at 4°C for 2 hours. Goat anti-rabbit and anti-mouse IgG antibodies conjugated with Alexa Fluor 594 and Alexa Fluor 647, respectively, were used (AB150080 and AB48389, respectively, Abcam, Inc.). The sample was mounted on a glass slide in Vectashield medium (Vector Laboratories, Inc.), and then imaged by confocal microscopy with Leica TCS SP8 DLS confocal microscope using an oil-immersion objective lens (HC PL APO CS2 40× 1.3NA, Leica).

### Image analysis

Image processing was done using ImageJ (27) and MATLAB (MathWorks, Inc.). Mosaics were created using an ImageJ stitching plugin (28). SHG normalization and thickness estimation was done as previously described (10). Briefly, the composite SHG intensity was corrected for topography and normalized by dividing with the Fano factor. The thickness of the retinal nerve fiber was evaluated by means of context-free image segmentation, which was done through edge detection and histogram-based thresholding. The thresholded z-stack images were sum-projected and then multiplied with the z-step (2 μm) to obtain the thickness of the retinal nerve fibers. The SHG density was obtained by dividing the normalized SHG intensity by thickness. The volume of the retinal nerve fibers was evaluated by integrating the thickness over the region.

### Statistical analysis

Statistical analysis was performed using R (29). The homogeneity of variance and normality were verified by inspecting the residual plots and the Q-Q plot. The collinearity of the explanatory variables was tested by the variance inflation factors. The IOP, volume, and mean SHG density were analyzed as the response variables with a multiple linear regression model. Three main effects of age, sex, and diet were considered. Of three possible interactions, the term between sex and diet was dropped on the ground that the effect of NAM is not known to be dependent on sex. Also, to aid model specification, the fit of multiple linear regression model was assessed by the adjusted R-squared. The alpha level for statistical significance was 0.05.

## Supporting Information

**Fig S1.**
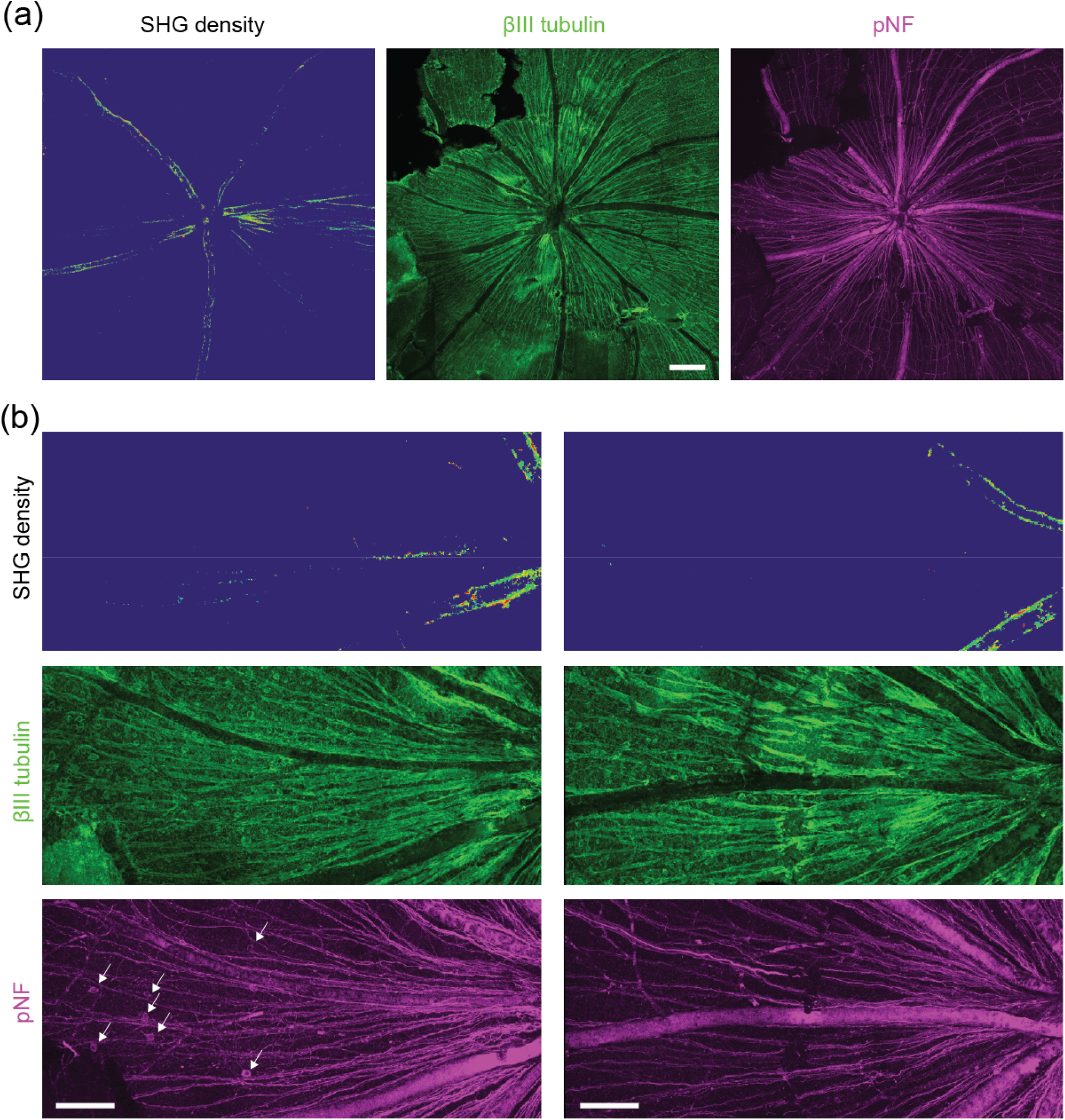
Microtubule deficit versus the distribution of cytoskeletal proteins, βIII tubulin and pNF. (a) Mosaics of a DBA retina (12 months age, female) by SHG and immunofluorescence against the cytoskeletal elements. Scale bars, 200 μm. (b) Comparing two sectors in different stages of RGC degeneration. Arrows, pNF+ RGC somas. Scale bars, 100 μm.

## References

1. Quigley HA, Broman AT. The number of people with glaucoma worldwide in 2010 and 2020. British Journal of Ophthalmology. 2006;90(3):262–7.

2. Tham YC, Li X, Wong TY, Quigley HA, Aung T, Cheng CY. Global prevalence of glaucoma and projections of glaucoma burden through 2040 A Systematic review and meta-analysis. Ophthalmology. 2014;121(11):2081–90.

3. Huang XR, Knighton RW. Microtubules contribute to the birefringence of the retinal nerve fiber layer. Investigative Ophthalmology & Visual Science. 2005;46(12):4588–93.

4. Huang XR, Knighton RW, Cavuoto LN. Microtubule contribution to the reflectance of the retinal nerve fiber layer. Investigative Ophthalmology & Visual Science. 2006;47(12):5363–7.

5. Fortune B, Burgoyne CF, Cull G, Reynaud J, Wang L. Onset and progression of peripapillary retinal nerve fiber layer (RNFL) retardance changes occur earlier than RNFL thickness changes in experimental glaucoma. Investigative Ophthalmology & Visual Science. 2013;54(8):5653–60.

6. John SWM, Smith RS, Savinova OV, Hawes NL, Chang B, Turnbull D, et al. Essential iris atrophy, pigment dispersion, and glaucoma in DBA/2J mice. Investigative Ophthalmology & Visual Science. 1998;39(6):951–62.

7. Danias J, Lee KC, Zamora MF, Chen B, Shen F, Filippopoulos T, et al. Quantitative analysis of retinal ganglion cell (RGC) loss in aging DBA/2NNia glaucomatous mice: Comparison with RGC loss in aging C57/BL6 mice. Investigative Ophthalmology & Visual Science. 2003;44(12):5151–62.

8. Libby RT, Anderson MG, Pang IH, Robinson ZH, Savinova OV, Cosma IM, et al. Inherited glaucoma in DBA/2J mice: Pertinent disease features for studying the neurodegeneration. Visual Neuroscience. 2005;22(5):637–48.

9. Inman DM, Sappington RM, Horner PJ, Calkins DJ. Quantitative correlation of optic nerve pathology with ocular pressure and corneal thickness in the DBA/2 mouse model of glaucoma. Investigative Ophthalmology & Visual Science. 2006;47(3):986–96.

10. Sharoukhov D, Bucinca-Cupallari F, Lim H. Microtubule imaging reveals cytoskeletal deficit predisposing the retinal ganglion cell axons to atrophy in DBA/2J. Investigative Ophthalmology & Visual Science. 2018;59(13):5292–300.

11. Wang JT, Medress ZA, Vargas ME, Barres BA. Local axonal protection by WldS as revealed by conditional regulation of protein stability. Proceedings of the National Academy of Sciences of the United States of America. 2015;112(33):10093–100.

12. Williams PA, Harder JM, Foxworth NE, Cochran KE, Philip VM, Porciatti V, et al. Vitamin B-3 modulates mitochondrial vulnerability and prevents glaucoma in aged mice. Science. 2017;355(6326):756–60.

13. Williams PA, Harder JM, Foxworth NE, Cardozo BH, Cochran KE, John SWM. Nicotinamide and WLDS act together to prevent neurodegeneration in glaucoma. Frontiers in Neuroscience. 2017;11.

14. Dombeck D, Kasischke K, Vishwasrao H, Ingelsson M, Hyman B, Webb W. Uniform polarity microtubule assemblies imaged in native brain tissue by second-harmonic generation microscopy. Proceedings of the National Academy of Sciences of the United States of America 2003;100(12):7081–6.

15. Lim H, Danias J. Label-free morphometry of retinal nerve fiber bundles by second-harmonic-generation microscopy. Optics Letters. 2012;37(12):2316–8.

16. Lim H, Danias J. Effect of axonal micro-tubules on the morphology of retinal nerve fibers studied by second-harmonic generation. Journal of Biomedical Optics. 2012;17:110502.

17. Howell GR, Libby RT, Marchant JK, Wilson LA, Cosma IM, Smith RS, et al. Absence of glaucoma in DBA/2J mice homozygous for wild-type versions of Gpnmb and Tyrp1. BMC Genetics. 2007;8:45.

18. Pease ME, Cone FE, Gelman S, Son JL, Quigley HA. Calibration of the TonoLab tonometer in mice with spontaneous or experimental glaucoma. Investigative Ophthalmology & Visual Science. 2011;52(2):858–64.

19. Dieterich DC, Trivedi N, Engelmann R, Gundelfinger ED, Gordon-Weeks PR, Kreutz MR. Partial regeneration and long-term survival of rat retinal ganglion cells after optic nerve crush is accompanied by altered expression, phosphorylation and distribution of cytoskeletal proteins. Eur J Neurosci. 2002;15(9):1433–43.

20. Soto I, Pease ME, Son JL, Shi XH, Quigley HA, Marsh-Armstrong N. Retinal ganglion cell loss in a rat ocular hypertension model Is sectorial and involves early optic nerve axon loss. Investigative Ophthalmology & Visual Science. 2011;52(1):434–41.

21. Pang IH, Clark AF. Inducible rodent models of glaucoma. Progress in Retinal and Eye Research. 2020;75.

22. Jakobs TC, Libby RT, Ben YX, John SWM, Masland RH. Retinal ganglion cell degeneration is topological but not cell type specific in DBA/2J mice. Journal of Cell Biology. 2005;171(2):313–25.

23. Schlamp CL, Li Y, Dietz JA, Janssen KT, Nickells RW. Progressive ganglion cell loss and optic nerve degeneration in DBA/2J mice is variable and asymmetric. BMC Neuroscience. 2006;7.

24. Howell GR, Libby RT, Jakobs TC, Smith RS, Phalan FC, Barter JW, et al. Axons of retinal ganglion cells are insulted in the optic nerve early in DBA/2J glaucoma. Journal of Cell Biology. 2007;179(7):1523–37.

25. Savinova OV, Sugiyama F, Martin JE, Tomarev SI, Paigen BJ, Smith RS, et al. Intraocular pressure in genetically distinct mice: an update and strain survey. Bmc Genetics. 2001;2.

26. Renier N, Wu ZH, Simon DJ, Yang J, Ariel P, Tessier-Lavigne M. iDISCO: A Simple, RapidMethod to Immunolabel Large Tissue Samples for Volume Imaging. Cell. 2014;159(4):896–910.

27. Schneider CA, Rasband WS, Eliceiri KW. NIH Image to ImageJ: 25 years of image analysis. Nature Methods. 2012;9(7):671–5.

28. Preibisch S, Saalfeld S, Tomancak P. Globally optimal stitching of tiled 3D microscopic image acquisitions. Bioinformatics. 2009;25(11):1463–5.

29. R Core Team, 2024. R: A Language and Environment for Statistical Computing, Vienna, Austria. Available at: https://www.R-project.org/.

